# Interferometric Fluorescent Cross Correlation Spectroscopy

**DOI:** 10.1101/671016

**Authors:** Ipsita Saha, Saveez Saffarian

**Affiliations:** Center for Cell and Genome Science, University of Utah; Department of Physics and Astronomy, University of Utah; Department of Biology, University of Utah

## Abstract

We present a method utilizing single photon interference and fluorescence correlation spectroscopy (FCS) to simultaneously measure transport of fluorescent molecules within aqueous samples. Our method, within seconds, measures transport in thousands of homogenous voxels (100 nm)^3^ and eliminates photo-physical artifacts associated with blinking of fluorescent molecules. A comprehensive theoretical framework is presented and validated by measuring transport of quantum dots, associated with VSV-G receptor along cellular membranes as well as within viscous gels.

## Main

As our understanding of cellular environments advances, the non-equilibrium, non-steady state nature of chemical reactions in biology becomes apparent^1-4^. Since these reactions are not at equilibrium, transport needs to be measured simultaneously across the sample to uncover correlations and complex relationships. Measurements of transport in microscopic systems and cells have advanced significantly using single particle tracking methods ^5, 6^ and with the application of high resolution localization techniques, it has been possible to track well defined molecules or molecular assemblies with nanometer precision within live cells ^7, 8^. Meanwhile simultaneously measuring diffusion and flow within the whole sample without identifiable traceable objects has remained out of reach of the particle tracking methods.

Diffusion and flow within the sample can be measured by analyzing the fluctuations in fluorescence due to underlying transport properties using Fluorescence Correlation Spectroscopy (FCS) ^9^. In recent years, FCS has advanced by introduction of cross correlation ^10^ as well as line scanning and pair correlation methods ^11, 12^. Unlike particle tracking, which requires resolving single particles, correlation spectroscopy can distill transport properties through fluctuations in a fluorescence signal corresponding to many molecules. Variations of FCS have been used to gather basic transport properties as well as connectivity maps of various compartments within live cells as reviewed in ^13^. The correlation spectroscopy methods are however limited by the low optical resolution along the optical axis as well as limited number of voxels analyzed during the experiments.

Interferometric fluorescence measurements were first introduced in interferometric PALM microscopy in 2009 ^14^. In these microscopes the photon wave-front interferes with itself with varying phase shifts and is imaged on multiple cameras. The result of using interference has been a significant increase in effective resolution of the optical microscope which now allows a routine ∼ (10*nm*)^3^ resolution for localizing single molecules within the sample ^15^.

Here we have merged Fluorescence Correlation Spectroscopy with interferometric single photon localizations to simultaneously measure transport properties in a cross section of the sample in 200 × 200 voxels with a voxel resolution of (100 *nm*)^3^. In our current setup, we can resolve transport along plasma membrane and or within viscous gels, however, due to limitations of detector speeds, we are unable to create a transport map of the cytosol of living cells. We discuss new detector technologies which should allow cytosolic measurements.

Our instrument is composed of two objectives focused on the sample from top and bottom as shown in the instrument diagram presented in Figure S1. In this geometry the sample is sandwiched between two coverslips and secured onto a micro positioning stage. The wave front emitted by the photon within the sample travels through two independent light paths initiated by the two objectives and is recombined in a three way prism with three phase variations of 0°, 120° and each focused on an independent cMOS camera para-focal with the sample plane. As long as the length of the two light paths remains within the coherence length of the emitted photon (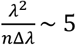 μm in which *λ* is the wavelength of the emitted fluorescence and *n* is the index of refraction) the photon would be detected by one of the cMOS cameras based on the interferometric probability of its detection. The point spread function of the scope is a convolution of typical optical microscope with an interferometric effect which is best described by a sine wave as defined below and experimentally verified in Figure S2:

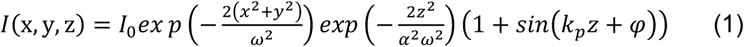

An experimental measurement of this point spread function is presented in Figure S2. Based on the point spread function, when a molecule travels by 80 nm along the optical axis its fluorescence would be detected on a different camera. During each exposure of the sample to the excitation light, which is provided into the sample through objective based TIRF, pixel images are acquired on all three cameras.

iFCCS works by calculating auto and cross correlation of the fluorescence signal registered on the three cameras as theoretically detailed in the supplementary materials. While in principle correlation functions can be calculated based on individual pixels, fluorescence from a single molecule is spread along an area of 3 × 3 pixels and therefore the fluorescence associated with each region of interest is calculated by summing the fluorescence within a mask of 3×3 pixels with the center pixel at the center of the region of interest, this is in agreement with the optimal pinhole size defined previously for FCS measurements ^16^. The background is calculated based on the average fluorescence from the 200×200 pixels as described in supplementary section and subtracted from the total fluorescence intensity calculated in each region of interest.

Experimental cross correlation functions are calculated by multiplying the signal from two ROI’s characterized by their center position (*x, y*) and phase *φ* of their corresponding cMOS chip:

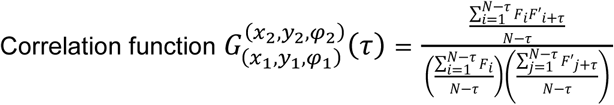

where 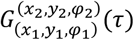 is the cross correlation function, *N* is the total number of frames, *F* is the integrated fluorescence minus background from the 9 pixel ROI at (*x*_1_, *y*_1_, *φ*_1_) and *F*′ is the integrated fluorescence minus background from the 9 pixel ROI at (*x*_1_, *y*_1_, *φ*_2_). The correlation functions were calculated using a multi-tau correlation algorithm ^17^.

One interesting feature previously described in PCF ^18^ is the prediction of an anti-correlation in various cross correlation situations. Here when the two detectors are 120° phase shifted and the molecule is in the plane of detector 1, interference will result in lowering of its registered signal on the other detectors. When the molecule diffuses along the optical axis, the signal strength in detector 2 increases while the signal in detector 1 decreases resulting in a detectable anti-correlation as shown in Figure S3. It is of note that, the cross correlation curves between the detectors, which are phase shifted by *φ* > 0, not only determine the diffusion and flow of the molecule along the optical axis with a very high resolution but also can distinguish between fluctuations arising due to actual physical movement of the molecule and other photo-physical activity of a fluorescent proteins. This is because when the molecule blinks or goes through any other photo-physical activity it affects both the detectors simultaneously. The characteristic anti-correlation between phase shifted cameras is a signature of molecular motion along the z-axis and therefore anti-correlation can only be detected during physical movement and not any other photophysical effect. Another distinct feature of this kind of correlation function is its asymmetry when flux is present in the system. When the system has pure diffusion the cross correlation functions are symmetric 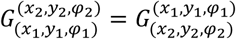 and when there is flux the cross correlations are asymmetric 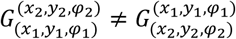. That is another advantage for determining the direction of flow of molecules in a particular observation volume.

We validated our method first using monte-carlo simulations. In our simulation 250 particles were placed in a 25×25×1*μm*^3^ volume with reflecting boundary conditions. Initially the particles were randomly distributed within the volume. The particles underwent Brownian diffusion with a diffusion coefficient of 10^−9^ *cm*^2^/*sec*. To determine the sensitivity of the method for detection of flow, molecules were subjected to directional flow (*v* = 0.4 *μm*/*sec*) within 6 regions of the sample as shown in Figure S4. Details of the simulation are provided within the supplementary. The calculated correlation functions and the simulated data are presented and explained in Figure 1. The correlation curves were fitted with the theoretical curves as derived in equation 3 from supplements and we obtained a diffusion coefficient of (0.89 ± 0.048) × 10^-9^ *cm*^2^/*sec* and flux of (0.36 ± 0.028) *μm*/*sec*.

**Figure 1:**
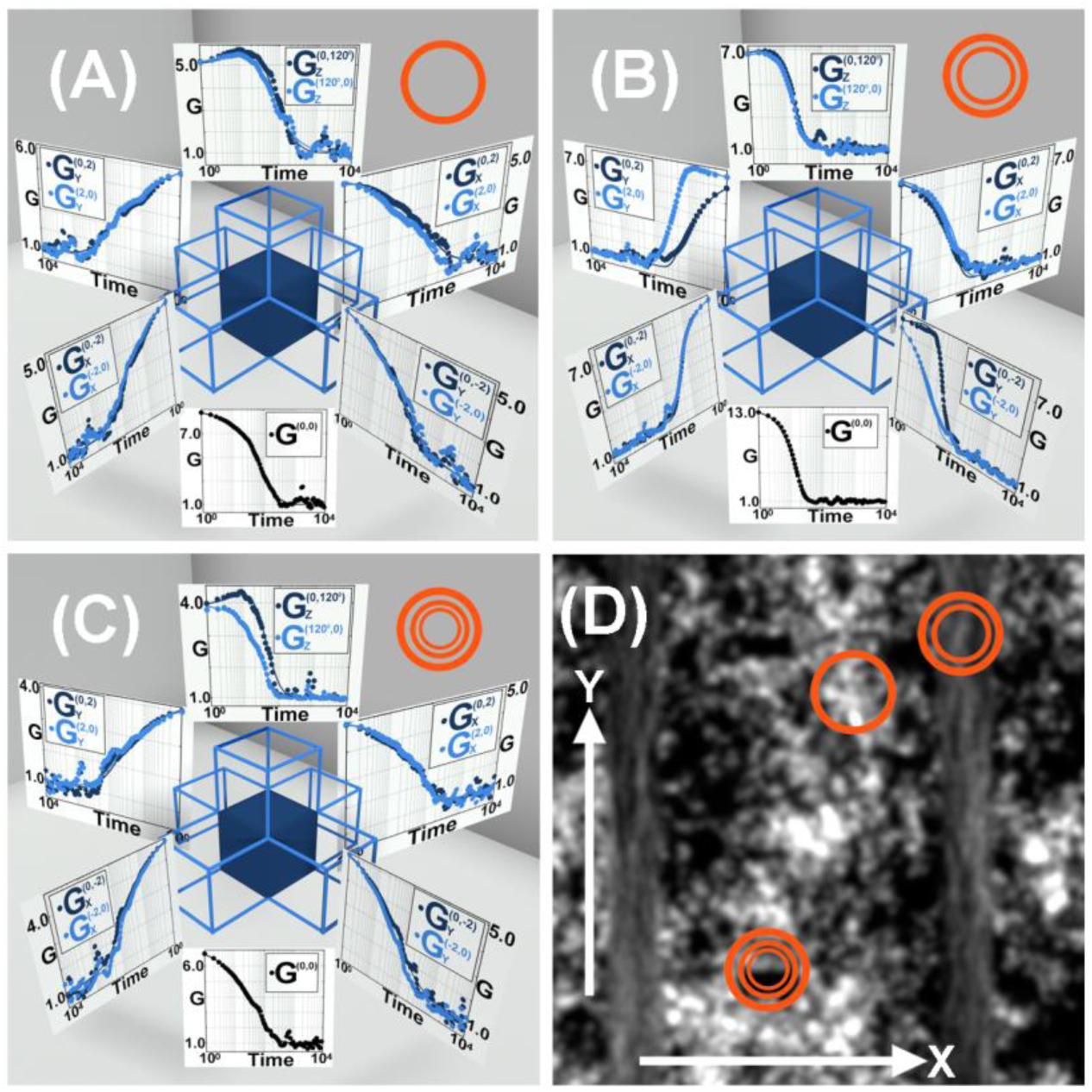
Transport measurements on particles in a box simulated by MonteCarlo dynamics. (A) shows the calculated correlation functions in a region where the particles undergo pure diffusion (*D* = 10^−9^ *cm*^2^/*sec*). (B) shows the correlation functions in a region where the particles were subjected to flow (*v* = 0.4 *μm*/*s*) along the Y axis. (C) shows the correlation functions in a region where the particles are subjected to flow along the Z axis. (D) shows the superimposed image from the simulated system for 20,000 frames with the regions marked which have been used for calculation in (A), (B) and (C). In each region, 1 auto correlation function, 8 cross correlation functions in the axial plane where the ROIs along the X and Y are separated by 2 pixels and 2 cross correlation along the optical direction between two 120° phase shifted images, has been calculated. In the legends the correlation functions has been represented as 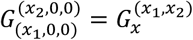, 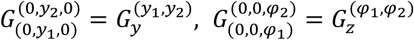 and 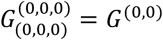. The fitted correlation curves gives a diffusion coefficient of (0.89 ± 0 048) × 10^−9^*cm*^2^/*sec* and flux of (0.36 ± 0 028) *μm*/*sec*.

To experimentally validate our method, a sample was prepared by adding Quantum dots (Quantum dots 605, Thermo Fisher Scientific) in 70% sucrose solution. The mixture was heated in a water bath at 80°*C* to dissolve crystals or air bubbles. This sample was sandwiched between two coverslips and was allowed to cool down to room temperature before imaging. The high viscosity of the sucrose created patches of inhomogeneity within the sample. 20,000 frames were acquired on three cameras with 1 *mSec* exposure and experimental cross correlation functions identical to the arrangement in figure 1 were calculated within the corresponding images. The superimposed image of the 20,000 frames is shown as well as correlation functions calculated within two selected ROIs as shown in Figure 2. While theoretically the sucrose solution should be homogeneous, we could detect directional flow within the ROIs as characterized by the asymmetry in the measured cross correlation curves shown in Figure 2. These flow vectors are likely due to temperature gradients within the sample. The 15–20 *nm* diameter quantum dots in 70% sucrose should theoretically yield a diffusion coefficient of ∼2 × 10^−9^*cm*^2^/*sec* at 33°*C* and we report to have obtained a diffusion coefficient of (2.68 ± 0.28) × 10^−9^*cm*^2^/*sec*.

**Figure 2:**
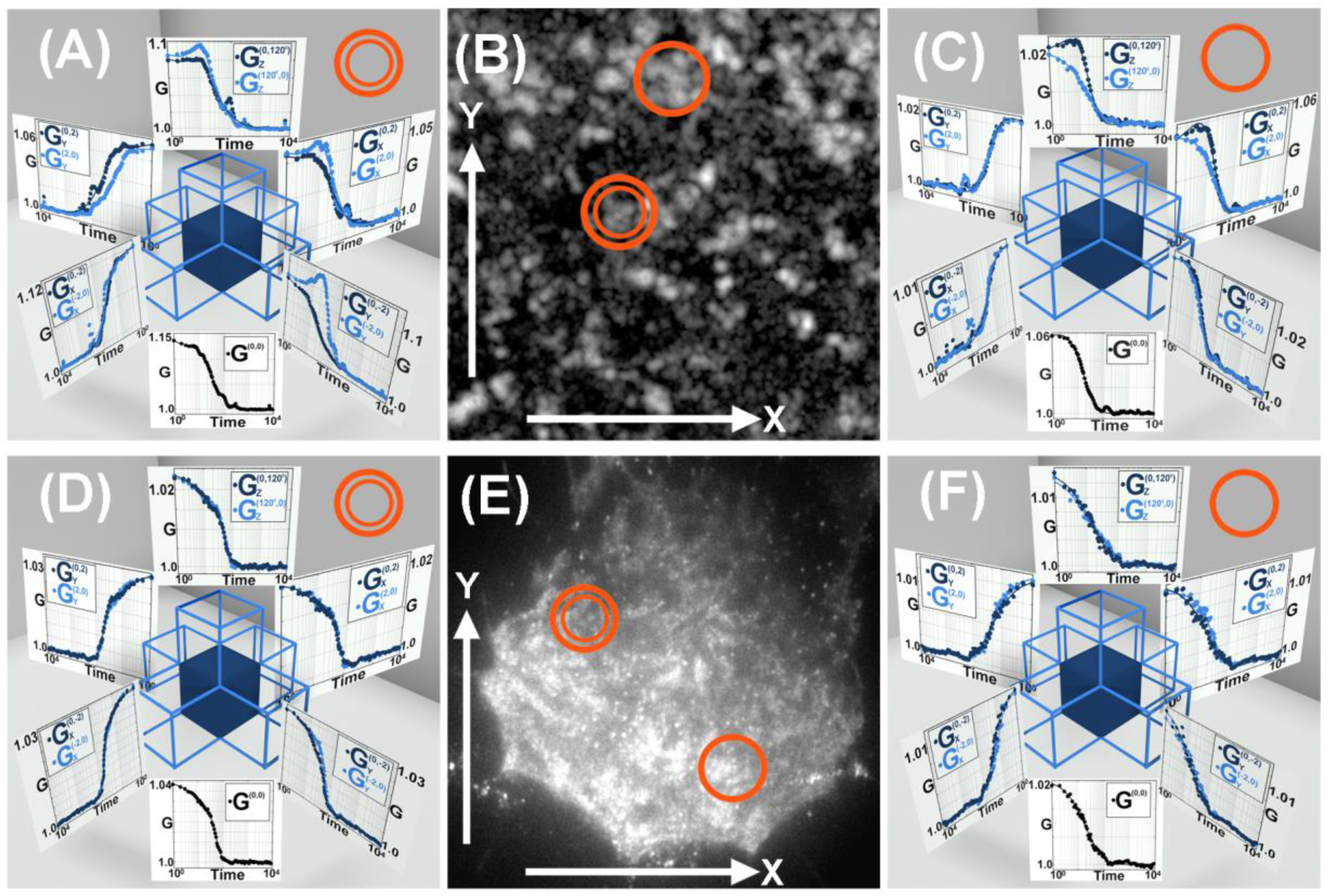
Transport measurements in a sample of quantum dots in sucrose solution and VSV-G diffusion on the membrane of HeLa cells. (A, B & C) show measurements of quantum dot transport within sucrose gel, (A&C) represent the correlation functions from two different regions of the sample. (B) is the superimposed image from the experimental dataset for 20,000 frames with regions marked which have been used for calculation in (A) and (C). The correlation functions have been fitted to obtain diffusion coefficient of (2.68 ± 0.28) × 10^-9^*cm*^2^/*sec* and regions of flow of 0.21*–*0.55 *μm*/*sec*. (D, E&F) represent measurments of transport of quantum dots associated with VSVG on the plasma membrane of HeLa cells. (D) and (E) represent the correlation functions from two different regions of the cell membrane. (E) is the image of the cell in which the correlation functions has been measured. The correlation functions has been fitted and we measured a diffusion coefficient of (1.28 ± 1.00) × 10^−10^*cm*^2^/*sec*. The correlation function calculation and representation is same as Figure 1.

We also measured pure 2*D* diffusion of VSV glycoproteins on cellular membranes. VSV-G was transfected in HeLa cells and incubated in 37°*C* for 12 *hours*. Biotinylated VSV-G antibody bound to streptavidin conjugate 605 quantum dots was added to these VSV-G transfected cells. This sample was then sandwiched after washing the cells so that the extra quantum dots are mostly removed from the solution. Thus the dynamics of the quantum dots reveals the diffusion of the VSV-G on the cell membrane. 20,000 frames were acquired on three cameras with 2 *mSeC* exposure and cross correlation functions were calculated as shown in Figure 2. These correlation functions reveal the 2D diffusion on the membrane and their symmetrical nature demonstrates absence of flow on the membrane of the cells. We report to have a diffusion coefficient of (1.28 ± 1.00) × 10^*-*10^*cm*^2^/*sec*.

Our experimental validation are currently limited by the speed of the cMOS cameras to 1 mSec per frame which limits the detection of diffusion to processes with diffusion coefficients below ∼3 × 10^−9^ *cm*^2^/*sec*. This limitation however can be overcome with faster detectors, one such possibility would be SPAD detectors recently developed for other high resolution applications ^19^ as well as SPAD detectors specialized for multi-tau correlation spectroscopy ^20^. SPAD detectors could potentially deliver frame rates within microseconds and therefore allow simultaneous measurements of diffusion and flow even within the cytosol of cells.

Cellular environments are crowded, compartmentalized and often host a few important enzymes fluxing through various states of their out of equilibrium cycle. It is fair to say that our understanding of the relationship of transport and chemical reactions in these systems is at its infancy, partly because we cant measure correlations between transport events in cellular environments in a large scale. The present iFCCS study, merges ideas in particle tracking and correlation spectroscopy with interferometric measurements and promises to create a transport map with homogenous voxels of the entirety of cellular environments.

## Acknowledgements

This study was supported by NIH R01GM125444 to (SS). The authors thank Rainer Daum for discussions and helpful comments during design of experiments.

## Author contributions

IS and SS designed the experiments, IS performed experiments and analyzed data, IS and SS wrote the manuscript.

## Supplementary Information

### Sample Prep

#### Quantum dots in sucrose

A 25 *mm* 1.5 Hestzig coverslip and an 18 *mm* 1.5 coverslip were used to sandwich the sample. The circular coverslips were thoroughly washed in 1 *M* sodium hydroxide solution followed by mQ water and then blow dried with nitrogen. 70% sucrose solution was prepared by dissolving the sucrose in 10 *mg*/*ml* casein solution in PBS. The sucrose solution was prepared by heating it in a water bath to 80°*C*. Care was taken not to insert any air bubble. 605 Quantum dots, by Thermo Fisher Scientific, were added to it when the sucrose completely dissolved and produced a clear viscous solution. The Quantum dots were heated with sucrose so the sample mixes well. After proper mixing, the sample was sandwiched between the coverslips and sealed with glue and allowed to cool down to room temperature. The casein prevented the quantum dots from sticking to the coverslips.

#### VSV-G HeLa cells

The coverslips were thoroughly washed in 1 *M* sodium hydroxide solution followed by mQ water and then blow dried with nitrogen and plasma cleaned. After cleaning, the coverslips were kept under UV irradiation in the biosafety cabinet for 2 hours before cells were plated. HeLa cells were plated on 25 *mm* #1.5 Hestzig coverslip. The cells were transfected with VSV-G GFP, 12 hours prior to experiment. 2 *µl* of biotinylated VSV-G antibody, by abcam, and 2 *µl* of streptavidin conjugate 605 Quantum dots, by Thermo Fisher Scientific, were mixed in 20 *µl* of *CO*_2_ independent media and incubated for 5 *Min*. Then this mixture was diluted in 180 *µl* of the media and added to the cells and incubated for 10 *Min*. The cells were then washed with the media and then sandwiched with 25 *mm* regular coverslips and vacuum grease.

#### Instrument specification and experimental details

The instrument is a prototype setup from Thermo Fisher Scientific as schematically described in Fig. S1. The sample was placed between two Nikon 60X Apo TIRF objectives of NA 1.49 and was illuminated by a 315 *mW* 561 *nm* laser. The 100 *nm* gold beads on the Hestzig slides were used to focus and calibrate the whole system. The custom 3-way beam splitter was adjusted so as to get the interference and 120°phase shift between the cameras (Hamamatsu Orca Flash 4.0 sCMOS) were obtained as seen in the calibration curves in Fig. S2. Data was collected on a 200 × 200 pixel area with 1 *msec* of exposure for 20,000 frames.

#### Simulation

The conditions for the simulated system were kept close to the real experiments. 250 particles were taken in a cube, which had a volume of 25 × 25 × 1 *μm*^3^. The pixel size in the simulation was chosen to be 100*nm*. The initial position of the particles was generated from rand function of MATLAB. The system had reflecting boundary condition. Spatial inhomogeneity was created where the particles were undergoing pure diffusional motion in some region while there were regions where the particles were subjected to directional flow as shown in figure S4. Areas of inhomogeneity were created in a way such that steady state condition was maintained in the system. A normalized random number generator (normrnd) determined the step sizes of each particle, the mean of which depended on our flux vector (0.4 *μm*/*sec*) and the standard deviation were determined by the diffusion coefficient (10^−9^*cm*^2^/*sec*). For the PSF, typical *ω* was chosen to be 264.5 *nm* and *α* was 4. The particles were excited by a 561 *nm* laser with a field depth of 300 *nm* and TIRF imaging conditions were maintained. Two detectors 120° phase shifted detected signals from these particles in the mentioned conditions for 20,000 frames with 1 *msec* of exposure. A poissonian random number determined the signals in the 200 × 200 pixel area with mean given by the PSF function as described in eq 1. The simulation code was written in MATLAB and run on the compute nodes with two Intel Xeon Gold 6130 CPUs, 32 CPU cores and 96 GB of RAM per node.

#### Mathematical details

The point spread function of the scope is a convolution of typical optical microscope with an interferometric effect which is best described by a sine wave as defined below and experimentally verified in Figure S2:

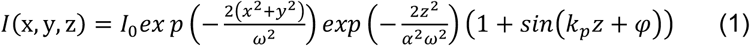

In which *ω* is the radial distance over which the intensity drops by 1/*e*^2^ and *αω* defines the axial distance over which the intensity drops by 1/*e*^2^ and *k*_*p*_ is the phase factor that is governed by the wavelength of excitation and numerical aperture of the objectives. The *φ* value is the interferometric phase shift of each camera and in our system it is either: 0°, 120° and 240°.

The probability of finding a molecule at any position is governed by the Smoluchowski equation:

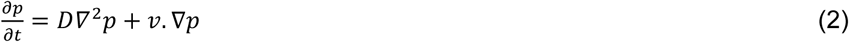

In which *D* is the diffusion coefficient and *ν* is the flux vector of the molecule.

Fluorescence Correlation functions are calculated based on probability density functions convoluted with the interferometric PSF function as defined in equation (1) and the total internal reflection excitation intensity profile *I*_*excitation*_ = *e*^−*z*/*d*^ where D is the TIRF penetration depth. Following the derivations as explicitly demonstrated in the supplementary, the generalized cross correlation function between the fluorescence detected from position (*x*_1_, *y*_1_, *φ*_1_) and (*x*_1_, *y*_1_, *φ*_2_) is calculated as:

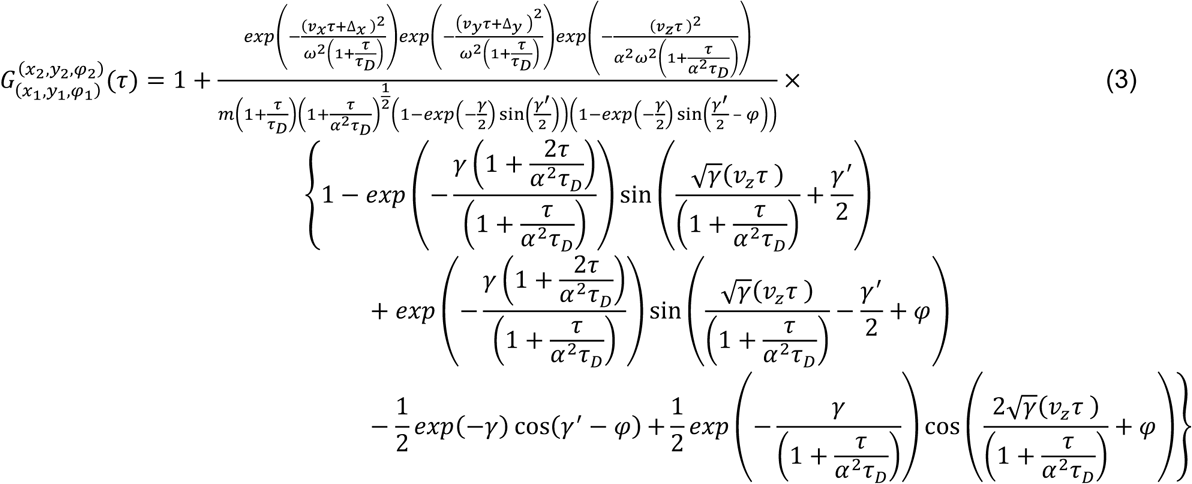

In which: 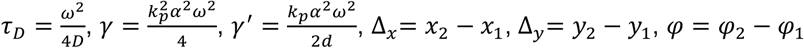, and *m* is the constant depending on the concentration of molecules in the observation volume.

For a system of volume *V*, the probability of the molecule being at (*x*′, *y*′, *z*′)at *t* = 0,

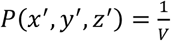

The fluorescence signal received by a detector (detector 1) from this molecule

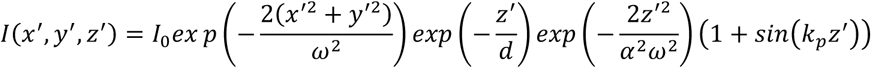

Since the molecule is has both diffusion as well as a directional flow, the probability of finding the molecule at (*x, y, z*), at *t* = *τ* can be found by solving Smoluchowski equation,

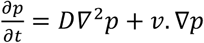

where *D* is diffusion coefficient and v is flux vector. Thus, the probability distribution function is

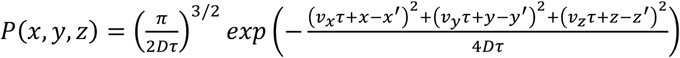

Due to the flux the fluorescence signal received at (*x* + Δ_*x*_, *y* + Δ_*y*_, *z*) from the same molecule by a detector which is phase shifted by *φ* from detector 1 is:

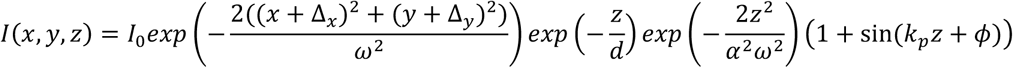

The generalized correlation function can be calculated as:

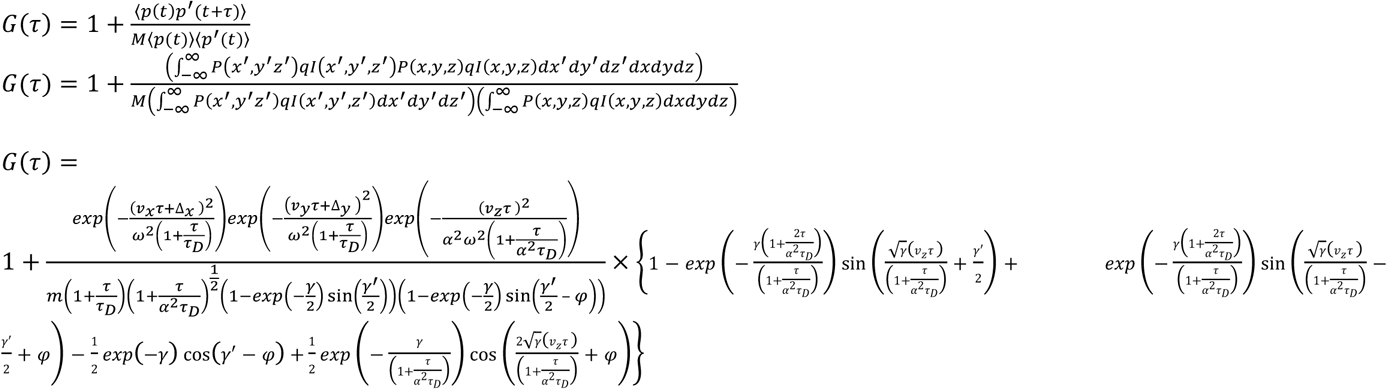

#### Calculation of fluorescence signal

For experimental correlation function the fluorescence signal is calculated by taking the integrated signal from the *m* × *m* pixel mask from which the background is subtracted from the entire *N* × *N* pixel that was imaged.

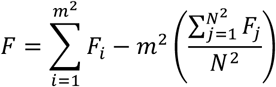

where *F*_*i*_ is the fluorescence signal from the pixels in the region of interest and *F*_*j*_ is the fluorescence signal from the pixels in the entire image.

---

**Figure S1:**
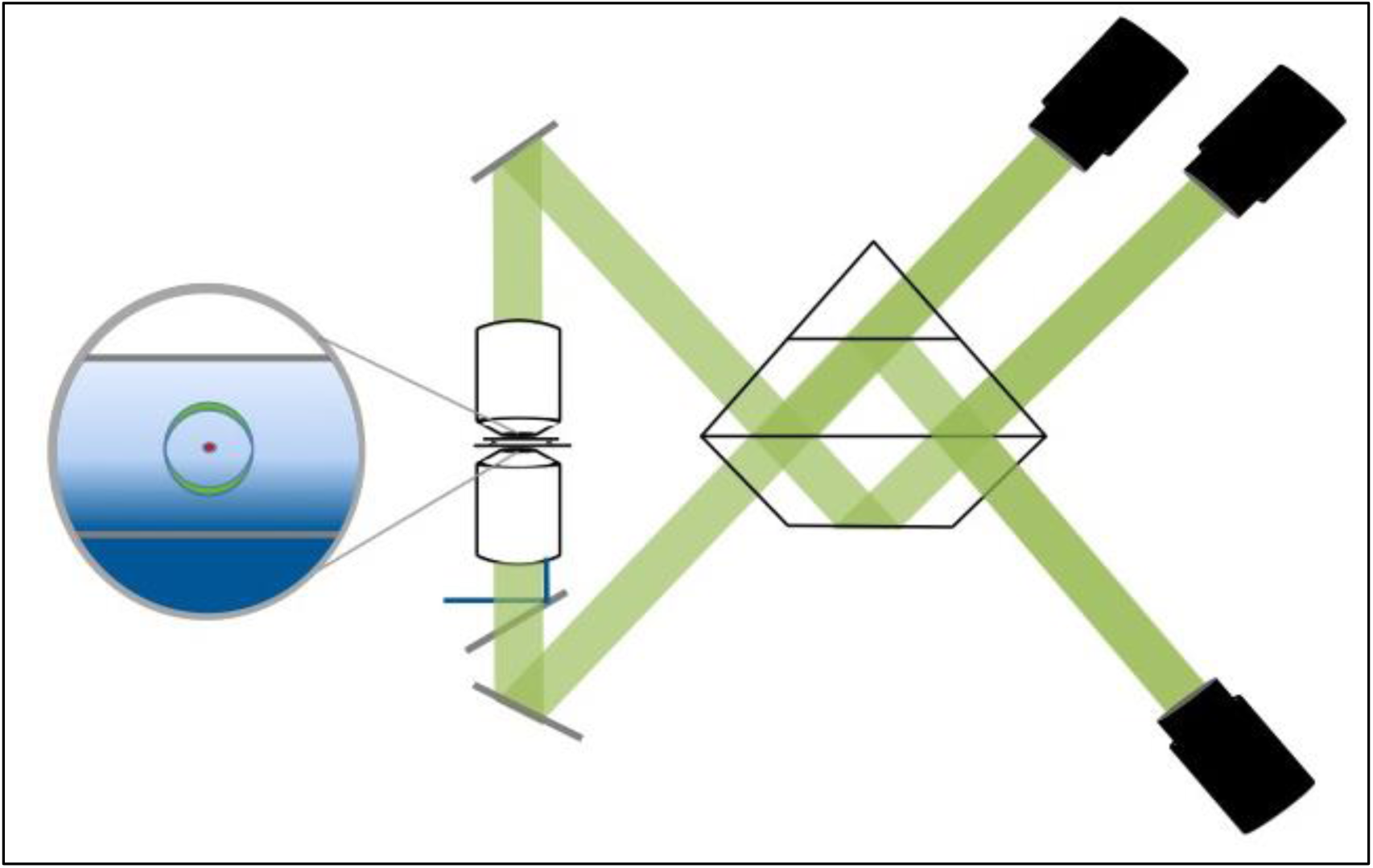
The experimental setup. Blue line demonstrate the path of the excitation TIRF laser, a single molecule is shown in red with a photon wave (green) emanating from the molecule. The green lines show the path of fluorescence through the instrument and prism until it reaches the three cameras. The instrument is similar to Steghel et al, except the camera’s are cMOS cameras.

**Figure S2:**
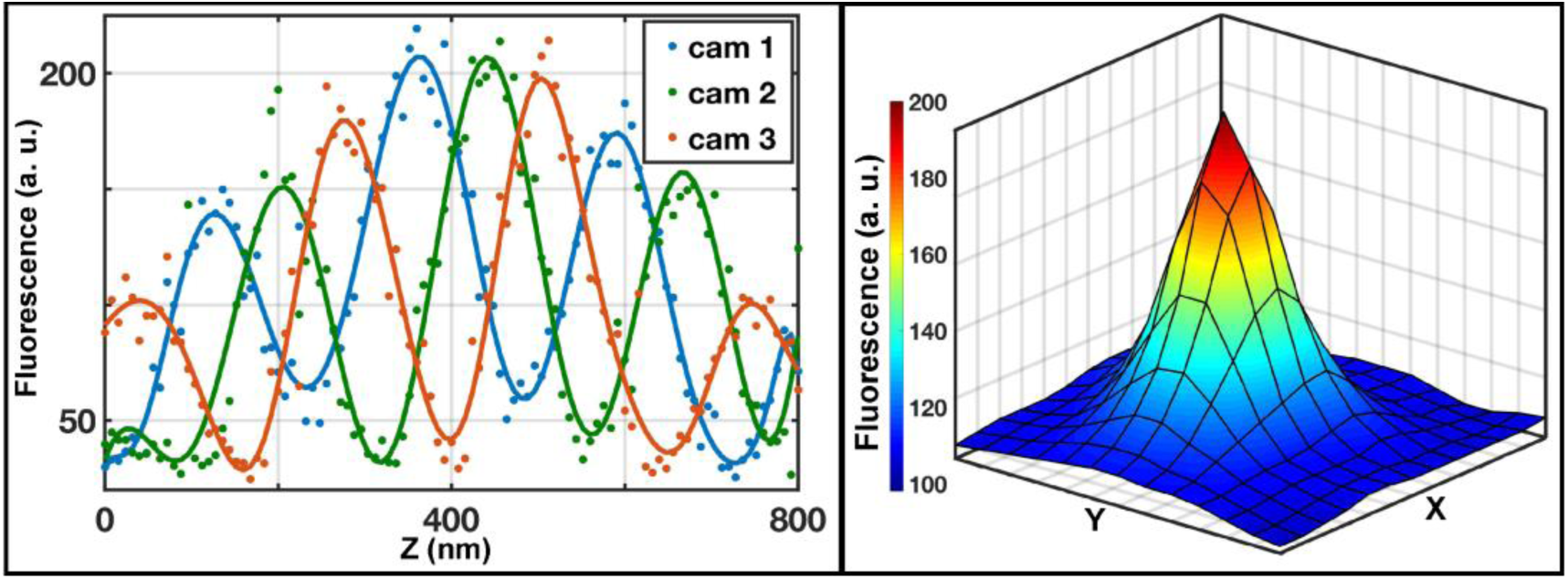
Calibration curve showing the interference effect and phase difference between the cameras. Calibration was performed by moving the sample in between the two objectives, in steps of 8*nm* for 101 planes, along the axial direction. At each plane the 3 cameras collected fluorescence signal from a fiducial.

**Figure S3:**
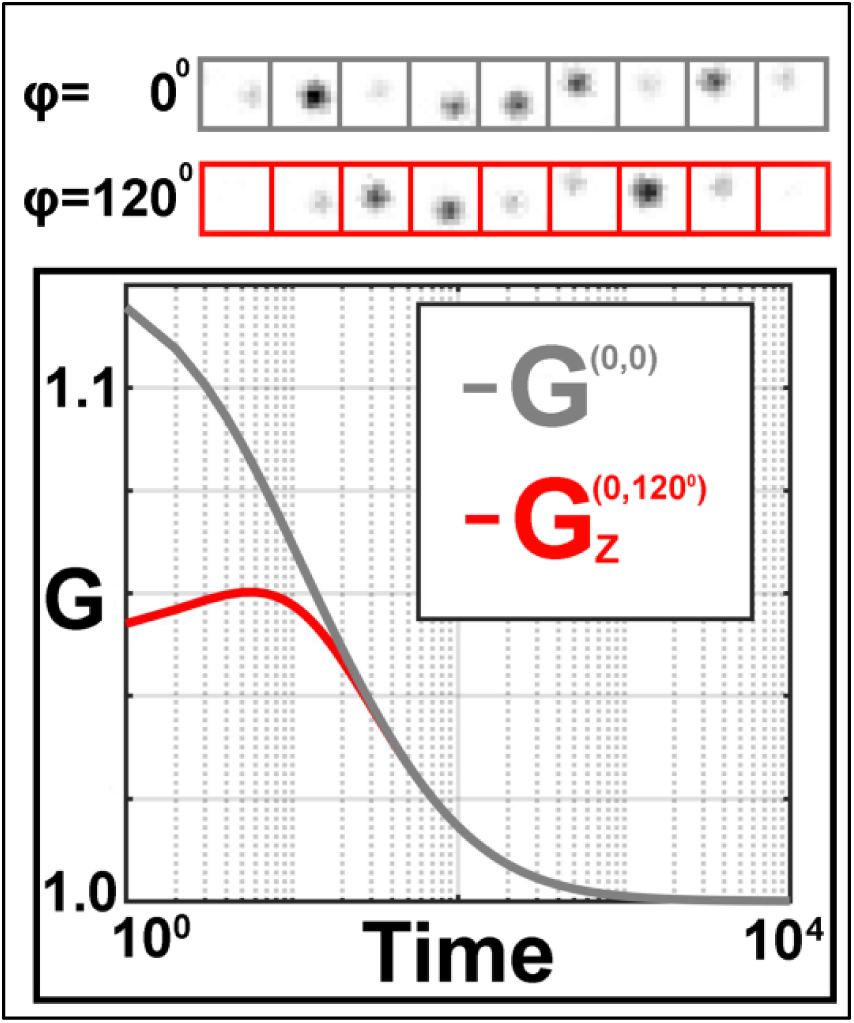
Demonstrates the movement of the molecule between the cameras and their associated correlation functions.

**Figure S4:**
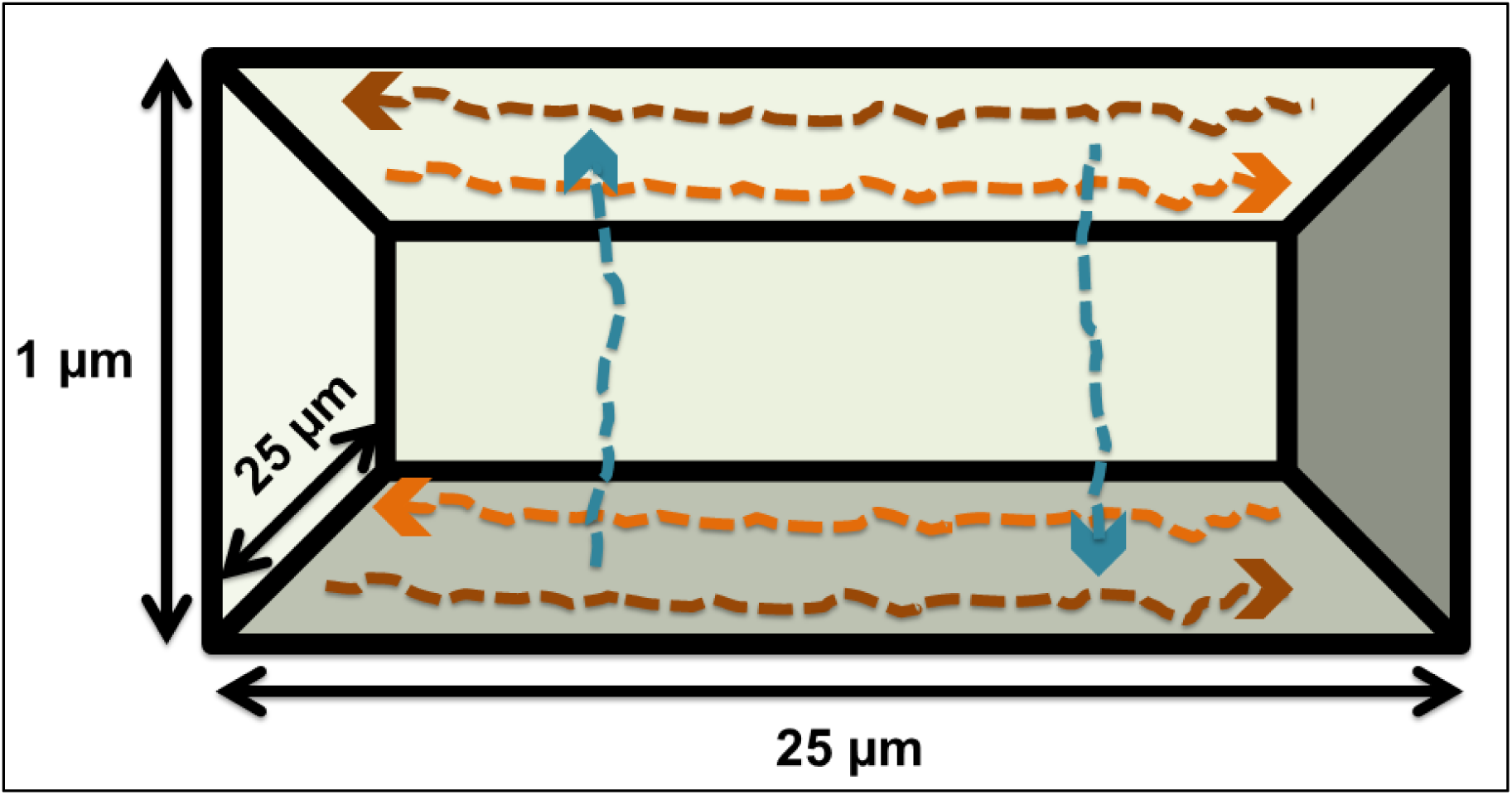
Represents the simulated system where the regions where the particles were subjected to flow are marked with arrows. The flow regions along the axial plane have a volume of 2×25×0.3 *μm*^3^ and the ones along the optical axis have a volume of 2×2×1 *μm*^3^.

